# Genomic analysis of *Klebsiella pneumoniae* causing community-acquired respiratory deaths among Zambian infants and children using targeted RNA-probe hybridization-capture metagenomics

**DOI:** 10.64898/2026.02.02.703236

**Authors:** Kenneth Lindstedt, Alyse Wheelock, Mulemba Samutela, Wasifa Kabir, Mwelwa Chasaya, Natasha Namuziya, Emilia Jumbe Marsden, Monica Kapasa, Chibamba Mumba, Bwalya Mulenga, Lisa Nkole, Rachel Pieciak, Victor Mudenda, Chilufya Chikoti, Benard Ngoma, Charles Chimoga, Sarah Chirwa, Lilian Pemba, Diana Nzara, James Lungu, Leah Forman, Edgar Simulundu, William MacLeod, Crispin Moyo, Somwe Wa Somwe, Kathryn E. Holt, Arnfinn Sundsfjord, Christopher J. Gill

**Author notes:** Corresponding author (technical questions): Kenneth Lindstedt, MBBS, PhD, Norwegian Centre for Detection of Antimicrobial Resistance, Department of Microbiology and Infection Control, University Hospital of North Norway, Norway., Corresponding author (study information): Alyse Wheelock, MD, Section of Preventive Medicine and Epidemiology, Chobanian & Avedisian School of Medicine, Boston University, Boston, MA, USA. Co-first authors, contributed equally (Kenneth Lindstedt, Alyse Wheelock).

## Abstract

*Klebsiella pneumoniae* (Kp) is a leading cause of neonatal and infant deaths in sub-Saharan Africa and frequently associated with antimicrobial resistance. Previously, we identified Kp as a major cause of fatal community-associated lower respiratory infections among infants and children under five years in Lusaka, Zambia, using postmortem tissue sampling and pathogen specific multiplex qPCR. In this follow-up study, we employed a novel culture-independent RNA-probe hybridization-capture metagenomic sequencing approach, targeting Kp pan-genome core and accessory genes, to perform in-depth genomic analysis of Kp from eleven post-mortem lung biopsy samples from seven of these children. Analysis detected Kp in all cases except one, which identified *Klebsiella quasipneumoniae* subspecies *similipneumoniae*. Core-genome multi-locus sequence typing (cgMLST) revealed six clonal groups (CG607, CG1123, CG10072, CG280, CG3648, and CG10344) belonging to five sublineages (SL607, SL17, SL280, SL37, and SL10072), with perfect concordance between paired samples from the same case. Two infants sampled the same month harbored SL607 lineages sharing 621 out of 629 cgMLST alleles, suggesting clonal spread. Kp capsule (K) loci were detected in all but one case and included potential vaccine targets KL25, KL23, and KL122. Antimicrobial resistance genes were widespread among samples, particularly encoding resistance toward aminoglycosides, β-lactams, sulphonamides, tetracyclines, and trimethoprim. Extended spectrum β-lactamases were identified in four cases, three of which were *bla*_CTX-M-15_. The acquired Kp sideophore yersiniabactin (lineage *ybt14*) was identified in both cases associated with SL607, and the acquired siderophore aerobactin (lineage *iuc5*) was identified in one of these, suggesting possible convergence of antimicrobial resistance and hypervirulence. The detection of Kp with extensive antimicrobial resistance causing fatal community acquired pneumonia signals a deeply concerning epidemiologic shift from a largely nosocomial pathogen. This calls for urgent epidemiological investigations to better understand the burden, transmission dynamics, antimicrobial resistances, and potential vaccine targets for Kp in other community settings across sub-Saharan Africa.

**Author Summary:** *Klebsiella pneumoniae* is a major cause of infections and death among newborns and young children, particularly in low-income countries, where it is frequently resistant to antibiotics. While well-known as a hospital-associated pathogen, we previously showed *K. pneumoniae* is also a leading cause of fatal community lung infections among infants and children in Lusaka, Zambia. In this follow-on analysis, we performed deeper genetic analysis of *K. pneumoniae* detected from the cluster of community pneumonia deaths using lung tissue samples from seven of these children. Since traditional bacterial cultures were unavailable, we instead used a novel approach that enriched and sequenced specific regions of the *K. pneumoniae* genome directly from the biopsy samples without culturing bacterial isolates. We identified five different *K. pneumoniae* genetic subtypes, known as sublineages. Two sublineages, which came from children sampled the same month, were highly similar, suggesting clonal spread. Multiple acquired antimicrobial resistance genes were detected across all sublineages. Acquired virulence factors, which may cause more aggressive infections, were also detected in two cases. We also identified capsule types previously suggested as potential vaccine targets. This study underscores the urgent need to better understand and address the emerging burden of antibiotic-resistant *K. pneumoniae* pneumonia and other invasive infections among infants and children in community settings in sub-Saharan Africa.

## Introduction

Historically, *Klebsiella pneumoniae* (Kp) has not been recognized as a significant cause of community-acquired acute lower respiratory infection (ALRI) among children, as informed by landmark epidemiologic research conducted in high-resource settings (1, 2). Previous large, multi-center studies of ALRI etiologies in low- to- middle-income countries (LMICs) have systematically excluded Kp from their analyses (3, 4). However, prior research suggests that nasal carriage and subsequent ALRI from enteric Gram-negative bacilli such as Kp vary geographically, with these pathogens featuring more prominently as causes of ALRI in warmer and more humid climates (5–9). More recent data from the Drakenstein birth cohort in South Africa strongly implicate Kp as an important cause of childhood community-acquired ALRI in certain settings (10). Moreover, Kp has emerged as a frequent pathogenic cause of childhood mortality in the Child Health and Mortality Prevention Surveillance (CHAMPS) network across sub-Saharan Africa and south Asia (11–13), including in deaths due to pneumonia (14). While the majority (86%) of deaths investigated through the CHAMPS network occurred in-hospital, of 329 deaths occurring in the community, 47 (14%) featured Kp in the causal chain, raising concern for possible community-acquired infection. We previously reported on a post-mortem study investigating 106 community deaths among children under age five in Lusaka, Zambia. Using multiplex PCR testing of lung samples collected via minimally invasive tissue sampling (MITS) (15–17), we were concerned to identify a cluster of community-acquired ALRI caused by Kp in 13 of 49 (27%) respiratory-related deaths.

Kp as a potential cause of childhood ALRI in community settings is alarming because 1) this pathogen is not well-covered by current treatment guidelines (18) and 2) global molecular epidemiology of Kp has been characterized by rising prevalence of multidrug resistant (MDR) and hypervirulent sublineages, as well as convergence of MDR and hypervirulent phenotypes (19). Molecular epidemiology data are sparser in sub-Saharan Africa than other regions of the world, but evidence indicates widespread MDR among clinical Kp isolates (13, 20–22). Genotypic and phenotypic studies have shown common to near-universal levels of extended-spectrum β-lactamase (ESBL) carriage among Kp in African settings, and molecular studies have revealed frequent detections of virulence factors, particularly the acquired siderophore yersiniabactin (23–27).

Previous in-depth genomic analysis has revealed Kp phylogeny is composed of a highly diverse population structure, consisting of hundreds of deep branching lineages (28). While acquisition of resistance and virulence determinants is widespread throughout the phylogeny, the majority of AMR and hypervirulence within Kp has been disproportionately driven by the expansion of a distinct subset of globally disseminated lineages, termed ‘global problem clones’ (28). Monitoring Kp genomic epidemiology is therefore of high importance to understand regional and international patterns of clonal spread, identify high-risk lineages, and inform infection control and intervention strategies, including vaccine development.

The finding of community deaths due to Kp ALRI in our original analysis was surprising and carried important clinical and public health implications. We therefore sought to better characterize the causative Kp in terms of antimicrobial resistance genes, virulence factors, lineage and capsule serotype by sequencing Kp DNA detected from the preserved lung tissue biopsies. Due to the unavailability of bacterial culture from these samples, we utilized culture-independent targeted hybridization-capture metagenomic sequencing to achieve these aims.

## Results

### Description of cases

Our previous post-mortem surveillance study enrolled 121 deceased infants and children between January 2020 to December 2022, with pauses in enrollment due to the COVID-19 pandemic. As previously reported, a total of 106 cases had complete molecular and histopathologic data and were adjudicated by the DeCoDe panel, of which 49 (46%) deaths were determined to be due to respiratory causes (15). Among these, Kp pneumonia was the most common respiratory cause of death, accounting for 13 (27%) of the respiratory deaths. For Kp targeted hybridization-capture analysis, DNA was re-extracted from 16 paired samples of 8 selected cases with the strongest molecular signal supporting Kp pneumonia. Of these, 5 samples proved insufficient for target hybridization-capture due to low DNA concentration (<10 ng/µL) and poor-quality metrics (260/280 ratio <1.5 or >2.2; 260/230 ratio <0.5 or >2.3). The remaining 11 samples from 7 deceased infants and children (4 paired samples and 3 individual samples) had sufficient quality and proceeded to target metagenomic hybridization-capture sequencing (**Supp. Table 1**).

The 7 deceased infants and children whose samples underwent Kp probe hybridization-capture analysis ranged in age from 7 days to 20 months and were enrolled in the study between February 2020 and December 2022. Based on enrollment criteria, none were hospitalized at the time of death; the deaths all occurred at home, en route to care, or while awaiting medical evaluation at a healthcare facility. All the deceased had a cough preceding their death, some had notable shortness of breath, and other symptoms reported included fever, vomiting, and diarrhea (**Table 1**). From the narratives provided by family members describing the events leading up to the death, some of the deceased had a noted prior interaction with a health facility, including prior hospitalizations (**Supp. Table 2**). In all but one case (where the information was insufficient), the DeCoDe panel determined that the death had been preventable, such as through earlier access to care, earlier escalation of care, avoiding inappropriate discharge, or via infection prevention measures during a prior hospitalization.

**Table 1:** Demographics of the deceased, case characteristics, *Klebsiella pneumoniae* sublineages and clonal groups detected among 7 infants and children who died of suspected *Klebsiella pneumoniae* acute lower respiratory tract infection in the community in Lusaka, Zambia. Kp, *Klebsiella pneumoniae*; Kqs, *Klebsiella quasipneumoniae* subsp. *similipneumoniae*.

| Case | Age | Sex | Month and<br>year | Location of death | Cough/<br>severe cough | Non-respiratory | Species | Sub<br>lineage | Clonal<br>Group |
| --- | --- | --- | --- | --- | --- | --- | --- | --- | --- |

|  |  |  | of death |  | (duration<br>in days) | symptoms |  |  |  |
| --- | --- | --- | --- | --- | --- | --- | --- | --- | --- |
| 2337 | 2 weeks | Female | Dec 2022 | En route to a facility | Yes (1) | Fever | Kp | SL607 | CG607 |
| 4803 | 7 months | Male | Dec 2022 | At home | Yes (2) | Fever,<br>vomiting,<br>diarrhea | Kp | SL607 | CG607 |
| 4771 | 8 months | Male | Oct 2022 | Waiting to be<br>seen by a doctor | Yes (2) | Vomiting,<br>diarrhea | Kqs | SL10072 | CG10344 |
| 4796 | 17 months | Male | Nov 2022 | At home | Yes (1) | Fever | Kp | SL17 | CG1123 |
| 4201 | 7 days | Female | Feb 2020 | En route to hospital<br>or other health facility | Yes (2) | Fever,<br>vomiting,<br>diarrhea | Kp | SL17 | CG10072 |
| 4220 | 18 weeks | Male | Jun 2020 | En route to hospital or<br>other health facility | Yes (7) | Fever | Kp | SL280 | CG280 |
| 4774 | 20 months | Male | Oct 2022 | Waiting to be seen<br>by a doctor | Yes (1) | Fever | Kp | SL37 | CG3648 |

Histopathology findings varied across participants as well as across the 6 biopsies obtained from each of the deceased, from interstitial expansion to histologic pneumonia. Most of the 7 individuals included in this study had Kp detected by FTD Respiratory Pathogens 33 multiplex PCR in 5 or 6 out of all 6 biopsies sent for molecular testing (**Supp. Table 3**). Most cases had other pathogens detected by the multiplex PCR on one or more biopsies; based on the totality of evidence (including number of detecting biopsies, qPCR cycle thresholds, host and pathogen characteristics), these were considered not to be the primary driver of the infection. Reanalysis of the 11 biopsy samples using the highly Kp-specific ZKIR-qPCR (29, 30) confirmed the presence of Kp in all samples, ranging from 45.5 to 6586.4 genome copies/ng DNA (median 147.6) (**Supp. Table 1**). A strong correlation was also observed between the ZKIR-qPCR quantified abundances and the FTD Respiratory Pathogens 33 multiplex PCR Kp Cq values (Pearson’s r = -0.94, *p* < 0.001, **Supp. Fig 1**), demonstrating high concordance between these two qPCR assays.

### Hybridization-capture of *K. pneumoniae* by targeted RNA-probe metagenomics

We performed in-depth genomic analysis of Kp lineages from lung biopsy samples by employing RNA-probe hybridization-capture metagenomics targeting Kp core and accessory genes (see Methods). This targeted Kp hybridization-capture was able to overcome considerable residual human DNA contamination.

Short-read sequencing of the 11 target-enriched samples produced an average of 30.3 x 10^6^ 75 bp paired-end reads per sample (range: 5.7 x10^6^ – 51.2 x10^6^). Following removal of human DNA, an average of 2.4 x10^6^ bacterial reads per sample remained (range: 6.4 x10^4^ - 13.5 x10^6^), indicating considerable residual human DNA contamination in samples despite targeted Kp hybridization-capture. Of these bacterial reads, an average of 57.8% were assigned to Kp (range: 30.1% - 89.6%), with the number of on-target Kp reads correlating strongly with the abundance of Kp in samples pre-hybridization-capture (Pearson’s r = 0.91, p < 0.001, **Supp. Fig 2**) (**Supp. Table 4**). With these limitations noted, our analysis was able to detect key genomic features of the Kp lineages, including sublineage (SL), clonal group (CG), capsule serotype, as well as acquired AMR genes and virulence determinants (**Table 2**).

**Table 2.**
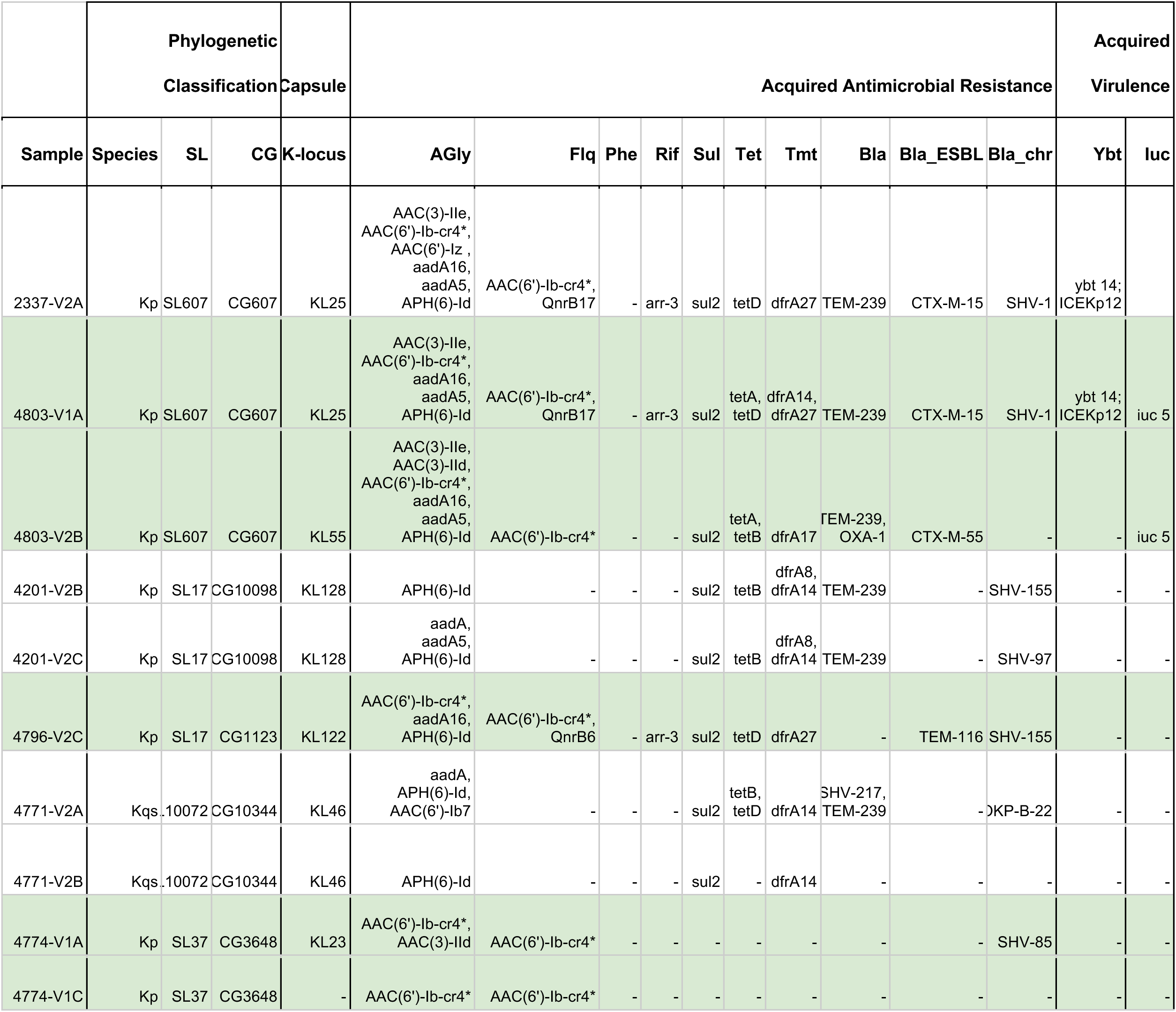
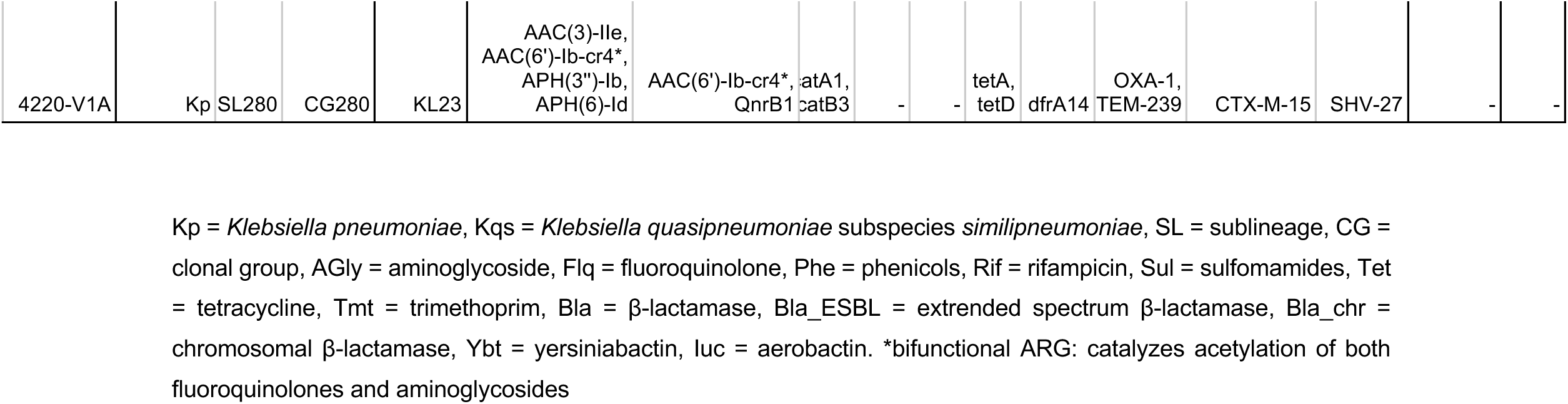
Key genomic features of *Klebsiella pneumoniae* sequenced from post-mortem lung biopsies from 7 infants and children who died of suspected *Klebsiella pneumoniae* acute lower respiratory infection in Lusaka, Zambia.

### Determination of *K. pneumoniae* cgMLST and MLST from enriched samples

Targeted RNA-probe metagenomics allowed for sequence typing according to current conventions. SL and CG analysis of Kp from the target enriched samples was performed using the previously described 629-gene core genome multilocus sequence typing (cgMLST) scheme and dual bar-coding system (31, 32). A median of 627 out of 629 cgMLST genes were detected per sample (IQR 625.5 - 628.5) (**Supp. Table 5**). When profiled against the Kp cgMLST profile database available through BIGSdb (https://bigsdb.pasteur.fr/klebsiella/), five SLs and six CGs were identified from the 11 enriched metagenomic samples: SL607 (CG607), SL17 (CG10098), SL17 (CG1123), SL37 (CG3648), SL10072 (CG10344), and SL280 (CG280) (**Table 1**). cgMLST profiles from enriched samples matched to cgMLST profiles within the database between 159 to 615 out of the 629 allelic positions (median 564) (**Supp. Table 5**). A strong correlation was observed between the number of matching alleles and on-target Kp reads in samples post-hybridization-capture (Spearman’s rho = 0.95, *p* < 0.001) (**Supp. Fig 3**), suggesting this was a major factor determining correct allele detection. All paired samples from the same case had identical SLs and CGs identified, supporting accurate cgMLST reconstruction. Additionally, all identified SLs corresponded to *K. pneumoniae sensu stricto*, except SL10072 (case 4771), which was identified as *Klebsiella quasipneumoniae* subspecies *similipneumoniae* (Kqs). Notably, the SL607 cgMLST allele profiles detected in samples 4803-V1A and 2237-V2A, which were taken during the same month from different individuals, were perfectly concordant at 621 out of 629 detected cgMLST allelic positions, indicating a very high degree of phylogenetic relatedness between the Kp strains present in these two samples (**Supp. Table 6**).

No cgMLST genes were detected in the negative control sample above our cut-off limit (≥80% gene coverage with ≥90% sequence identity). This confirmed no significant Kp DNA contamination of samples occurred during the hybridization-capture procedure, either through background laboratory contamination or through cross-contamination between samples, giving confidence in our cgMLST gene detection.

To complement our cgMLST analysis, reconstruction of sequence types (STs) using the previously described seven-gene Kp MLST scheme was also performed (33). Although this approach produced less consistent ST matches between sample pairs, the closest matching MLST allele profile was still concordant with our cgMLST analysis in 7 out of 11 samples (**Supp. Table 7**). Notably, 6 out of the 11 samples detected either 6 or 7 out of 7 MLST genes with 100% gene coverage with 100% sequence identity. All these alleles matched their expected MLST profiles positions based on our cgMLST analysis, except samples 4796-V2C and 4771-V1A which had single allele mismatches. Like our cgMLST analysis, the number of detected MLST alleles that matched with the expected MLST allele profile correlated strongly with the number of on-target Kp reads in samples post-hybridization-capture (Spearman’s rho = 0.86, *p* < 0.001, **Supp. Fig 4**). Moreover, all samples that produced over 800,000 on-target Kp reads following hybridization-capture produced an MLST profile matching that predicted by cgMLST analysis. Thus, although less precise than cgMLST reconstruction, MLST analysis was able to produce accurate ST profiles in the majority of samples, supporting our cgMLST analysis. The strong dependence on the number of on-target Kp reads indicated MLST analysis is best suited to samples that produce a higher yield of reads from the targeted hybridization-capture method.

### Capsule type (K-locus) prediction

Prediction of the polysaccharide capsule synthesis locus (K-locus), a key potential target for future development of Kp vaccines, was successfully performed using hybridization-capture. K-locus prediction was performed by aligning reads to alleles corresponding to the K locus marker genes *wzi*, *wzy*, and *wzx*, using allelic sequences from the Kaptive database (34). Using this approach, K-locus marker genes were detected in all samples except 4774-V1C (**Table 2**). The K-loci predicted from these marker genes were concordant between both paired samples from cases 4771 (SL10072, KL46) and 4201 (SL17, KL128), while unpaired samples 4220–V1A (SL280), 4796-V2C (SL17), and 2337-V2A (SL607) predicted K-loci KL23, KL122, and KL25, respectively. A mismatch occurred between paired samples from case 4803 (SL607), with sample 4803-V1A predicting KL25, in agreement with SL607 and KL25 detected in sample 2337-V2A, while 4803-V2B predicted KL55. Notably, samples 4803-V2B and 4774-V1C had the two lowest number of Kp-assigned reads of all samples, both less than 100,000 reads, which may suggest these were below the detection threshold for accurate K-locus detection by this method.

### Acquired antimicrobial resistance genes

We next analyzed samples for the presence of antimicrobial resistance genes (ARGs) known to be acquired by Kp, which constrain effective options for clinical treatment. A high prevalence of acquired ARGs was observed in all samples, with a median of 9 acquired ARGs from a median of 5 resistance classes detected per sample (**Table 2**). A strong concordance was observed in ARG profiles detected between paired samples from the same case. SL607 from samples 4803-V1A, 4803-V2B, and 2337-V2A, as well as SL280 from sample 4220-V1A, were associated with the highest number of acquired ARGs (n = 13, 13, 12, and 12, respectively). ARGs encoding acquired resistance to aminoglycosides, trimethoprim, β-lactams, tetracyclines, and sulphonamides were highly prevalent, detected in n = 11, 9, 8, 8, and 8 samples, respectively. Transferable fluoroquinolone resistance determinants were also detected in seven samples; *AAC(6’)-Ib-cr4* plus *Qnr* in four samples and *AAC(6’)-Ib-cr4* only in three samples.

Within the β-lactamase class, the globally disseminated extended-spectrum β-lactamase (ESBL), *bla*_CTX-M-15_, was detected in three samples; 2337-V2A and 4803-V1A associated with SL607, and 4220-V1A associated with SL280. Notably, the other sample from case 4803 associated with SL607 (4803-V2B) had a different CTX-M allele detected, *bla*_CTX-M-55_, possibly a misidentification due to low recovered reads within this sample. Another ESBL, *bla*_TEM-116_, was detected in sample 4796-V2C associated with SL17. The additional acquired broad-spectrum β-lactamases, *bla*_TEM-239_ and *bla*_OXA-1_, were detected in 7 and 2 samples, respectively.

The vast majority of Kp and Kqs lineages carry the intrinsic broad-spectrum β-lactamases *bla*_SHV_ and *bla*_OKP_, respectively, providing resistance to amoxicillin/ampicillin (35). This is of high clinical importance, as amoxicillin is the empiric therapy for outpatient community acquired pneumonia in Lusaka, Zambia (18). These β-lactamases were detected in all samples containing Kp (*bla*_SHV_) or Kqs (*bla*_OKP_) except 4774-V1C, 4803-V2B, and 4771-V2B. These samples had the lowest, second lowest, and fifth lowest number of on-target Kp reads recovered (median 62,020 reads), providing further support for a minimum threshold of approximately 100,000 reads for reliable gene detection. Thus, the Kp lineages associated with these cases of fatal community-acquired LRTIs were highly resistant, with most harboring multiple ARGs affecting clinically important drug classes.

### Acquired virulence determinants

Analysis of known acquired virulence loci amongst the identified Kp lineages revealed the presence of virulence factors in two cases. This analysis showed that Kp sublineage SL607 from samples 4803-V1A and 2337-V2A were associated with the acquired siderophore yersiniabactin (*ybt*), specifically lineage *ybt* 14 which is associated with the ICEKp12 mobile element (**Table 2**). Additionally, nine out of eleven alleles matching this *ybt* lineage were also detected in the other sample from case 4803 (4803-V2B) at ≥ 80% coverage and ≥ 99% sequence identity, highly suggestive of presence within this sample. None of these alleles, however, reached our minimum threshold of 50x coverage depth required for reliable detection. Similarly, samples 4774-V1A and 4774-V1C associated with SL37 had 8 and 6 of the 11 alleles matching the *ybt 9* lineage associated with ICEKp3, detected at ≥ 80% coverage and ≥ 95% sequence identity, suggestive of the presence of this locus within these samples, however these also did not meet our minimum threshold of coverage depth.

Both samples from case 4803 associated with SL607 also had the acquired siderophore aerobactin (*iuc*), specifically lineage *iuc5,* detected above the minimum identity and coverage thresholds. The other major known virulence loci which can be acquired by Kp (siderophore salmochelin, hypermucoidy locus *rmp*, and genotoxin colibactin) were not detected in any samples.

Detection of possible convergence of MDR and hypervirulence in sublineage SL607 in case 4803 by the acquisition of *iuc5* is a potentially important finding with significant potential public health implications. It was therefore important to investigate whether *iuc5* detected in this case may have been due to contamination with *Escherichia coli*, which is also a known carrier of this *iuc* lineage. Unique reads were assigned to *E. coli* in all 11 samples, ranging from 161 to 30896 (median: 13637, IQR: 3791 – 23003), suggesting either spurious hits during taxonomy profiling or low-level background contamination (**Supp. Table 8**). Notably, the two samples with *iuc5* detected, 4803-V1A and 4803-V2B, contained among the highest levels of *E. coli* reads detected, with 25012 and 22469 reads, respectively, ranking second and fourth highest overall. Thus, although detection of *iuc5* in case 4803 may suggest acquisition by Kp, resulting in a higher propensity to cause invasive infection, spurious detection from contaminating *E. coli* could not be ruled out.

### Plasmid content

Identification of multiple MDR-associated plasmid replicon markers by hybridization-capture provided insights into potential mechanisms of ARG acquisition and dissemination among Kp lineages. Analysis of plasmid content associated with Kp lineages identified seven different plasmid replicon types, with a median of four detected per sample (range 0 – 8) (**Supp. Table 9**). Detected plasmid replicon families included IncFIA, IncFIB, IncFII, IncI, IncN, IncR, and Col, with similar replicons detected between paired samples. Of these, the commonly Kp- and MDR-associated replicons Col(pHAD28), IncFIB(K), IncFIA(HI1), and IncR were most prevalent, detected in 9, 5, 5, and 5 samples, respectively. Other detected plasmid replicons associated with high-risk Kp clones and MDR carriage included IncFII(K), IncFII(pKP91), IncFII(pHN7A8), IncColRNAI, and IncN (28, 36, 37).

### Analysis of *K. pneumoniae* lineages from assembled contigs

To complement our read-based analysis, reads were assembled into longer contiguous sequences (contigs) and Kp lineages were analyzed for MLST, serotyping, acquired ARGs, and virulence determinants. Although contig-based analysis can have reduced detection sensitivity, reconstructing entire gene sequences can offer improved precision and confidence of results compared to read-based analysis alone. This analysis resulted in complete MLST profiles, containing perfect allele matches for all seven genes, in four samples (2337-V2A, 4803-V2B, 4796-V2C, and 4220-V1A) (**Supp. Table 10**). All these samples identified the same ST lineage as our read-based cgMLST and MLST analysis (ST607, ST607, ST17, and ST280, respectively). Of the remaining samples, all alleles with a 100% identical match to the MLST database produced the expected allele for that position based on our previous read-based analysis, except for one (*pgi* in sample 4771-V2B).

K-locus prediction from contigs was also highly concordant with our read-based analysis, with the best match K-locus agreeing with the K-locus detected from reads in almost all samples. Notably, the K-locus predicted in SL607 from case 4803 that had a mismatch between sample pairs from read-based analysis was concordant when performed on contigs. Both samples from this case predicted KL25, making it also concordant with SL607 detected in sample 2337-V2A. Additionally, paired samples 4774-V1A that detected KL23 and 4774-V1C which did not detect a K-locus from read-based analysis had a mismatch from contig-based analysis, predicting KL15 and KL51, respectively.

Analysis of acquired ARGs also produced highly concordant ARG profiles to our read-based analysis, with strong agreement between sample pairs. Similar to our read-based analysis, this demonstrated the presence of genes associated with resistance to β-lactams, aminoglycosides, trimethoprim, sulphonamides, and tetracycline was widespread among samples, as well as resistance to fluoroquinolones, MLS, and chloramphenicol in the majority of samples. Within the β-lactamase class, all ESBLs detected from contig analysis were concordant with our previous read-based analysis, with an additional detection of *bla*_CTX-M-14_ and *bla*_CTX-M-15_ in sample 4774-V1A which were undetected from reads.

Finally, analysis of acquired virulence from contigs also showed good agreement with read-based detection, with samples 2337-V2A and 4803-V1A associated with SL607 matching *ybt14* at 11 out of 11 allelic positions, while sample 4803-V1A also matched *iuc5* at four out of five allelic positions. Additionally, the acquired siderophore salmochelin (*iro*), lineage *iro5*, was also detected in sample 4803-V1A with two out of four allelic matches.

## Discussion

This analysis of a series of Kp pneumonia cases among children who died in the community in Lusaka, Zambia, revealed several previously identified Kp genetic lineages that had been observed in NICU settings and other environments. Notably, the sequencing revealed high rates of AMR, global problem clones SL17 and SL37 (36, 38), multiple virulence factors, and multiple capsular types. Two of the cases from the same month were both found to have SL607 sublineages that were highly phylogenetically similar, suggestive of clonal spread. Given that the majority of the deceased had no apparent contact with the hospital setting, the most parsimonious explanation for this cluster is that they represent spillover of hospital isolates that are now circulating outside the facility and causing community acquired infections. This has significant implications as it reinforces observations that the epidemiology of Kp has evolved and is no longer exclusively a hospital acquired pathogen. Given the lack of AMR testing, and the ubiquitous use of amoxicillin as first line therapy for pneumonia in community settings such as Lusaka, these findings should be viewed with significant alarm.

RNA-probe hybridization-capture metagenomic sequencing adds to evidence generated by our prior MITS study implicating Kp ALRI as the cause of deaths in these infants and children. While detection of Kp does not equate to invasive infection or cause of death, the histopathologic evidence and PCR data across multiple (six) samples adjudicated by an expert panel strongly supported Kp as the causative pathogen in these deaths. These findings, though limited by a small sample size, add to mounting evidence of a community burden of severe Kp, including as a cause of ALRI, in parts of sub-Saharan Africa and other LMIC settings. Of note, this study did not collect sufficient information to prove that the site of acquisition occurred in the community; narrative verbal autopsies did not mention any such hospital contact in the majority of cases.

If reproduced within larger studies, the implications of a shift in the epidemiology of Kp towards causing severe and fatal pediatric ALRI in the community should be viewed with extreme concern; this is especially true in resource-limited settings, where ALRI treatment is algorithmic and empiric, and almost never guided by pathogen diagnostics, let alone antimicrobial resistance testing. Current WHO guidelines for childhood pneumonia treatment at health facilities recommend amoxicillin for pneumonia without warning signs and ampicillin with gentamicin in severe pneumonia (18). Due to the chromosomally encoded broad-spectrum β-lactamase *bla*_SHV_, Kp is intrinsically resistant to amoxicillin/ampicillin (35). Among the 7 Kp ALRI deaths we investigated, aminoglycoside resistance genes were universal. Extended-spectrum β-lactamase, fluoroquinolone, and trimethoprim-sulphamethoxazole resistance determinants were also prevalent in our study. This leaves the carbapenem class, which is rarely available even in tertiary care settings in Lusaka, although notably, acquired resistance to this class has been detected recently within Kp isolated from Zambian hospitals (39, 40).

Several genomic findings bear highlighting. While most of the sublineages identified between children were distinct, two of the cases from the same month contained highly phylogenetically similar SL607 sublineages which shared 621 out of 629 cgMLST alleles, likely representing clonal spread. The biopsies associated with these related sublineages came from a neonate of 2 weeks and a 7-month-old infant, whose verbal autopsy described a hospital admission the previous week. Further information that could point to a common transmission source is not available. In contrast, the SL17 sublineages detected from two separate cases were less closely phylogenetically related, belonging to two different CGs, and thus were unlikely to represent a direct epidemiologic link.

Regarding the specific SLs identified in this study, SL607, SL17, SL10072, SL37, and SL280, each has been previously identified as known MDR-associated sublineages, and most have been reported to be important causes of hospital-acquired infections and outbreaks among neonates (27, 38, 41, 42). SL17 in particular is recognized as a MDR global problem clone that is prevalent throughout sub-Saharan Africa and has been reported as a cause of neonatal infections both within hospitals and the community (38, 43). SL37 is also a global problem clone that was identified among the most prominent Kp lineages responsible for neonatal sepsis in low- and middle-income countries within the BARNARDS study (23, 36). One detected sublineage, SL10072, was additionally found to belong to the species *K. quasipneumoniae subsp. similipneumoniae*, and encoded MDR. *K. quasipneumoniae* is increasingly recognized as a cause of invasive infection capable of acquiring MDR, including in sub-Saharan Africa (44, 45), supporting a potential need for increased surveillance of this species.

All SLs identified in this study, with the exception of SL10072, were also identified in a recent hospital-based study of Kp isolates from Malawi (27). Among these overlapping SLs, all except SL17 from case 4201 had matching K-loci with those predicted in our study and exhibited highly similar profiles of acquired ARGs. These findings suggest that the lineages identified in our cohort may form part of a broader circulating Kp population spanning both community and hospital settings across sub-Saharan Africa. Notably however, only a single sublineage (SL280) overlapped with a recent hospital-based study of Kp isolates from Zambia, which also had a matching K-locus (KL23) (46). While this limited overlap could indicate differences between community- and hospital-associated Kp populations within Zambia, it may also reflect temporal differences in sampling between studies, with several years separating sample collection resulting in shifts in the circulating lineage composition.

Another key finding was the detection of the acquired siderophores *ybt14* associated with SL607 in cases 2337 and 4803, as well as *iuc5* in case 4803. Analysis of all ST607 isolates available through PathogenWatch via Kleborate (n = 39, downloaded July 2025) (47) demonstrated that 66.7% of these were associated with carriage of *ybt14*, consistent with our findings. Notably, however, none of these isolates were associated with *iuc5* carriage. Acquisition of these siderophores may suggest convergence of MDR genes and hypervirulence factors within this lineage. Converged MDR-hypervirulent lineages may cause highly aggressive infections with limited treatment options that many primary and secondary health settings in limited-resource settings are unequipped to address. This finding therefore places the community death associated with this case within a well-documented and concerning trend of high mortality related to converged MDR-hypervirulent lineages of Kp, which is being observed globally, including other countries across sub-Saharan Africa (27, 48).

Serotyping of lineages causing community-acquired pneumonia may be an important contribution to choosing capsule targets for vaccine development, after further study in larger populations (49). Five different K-loci were identified within this study, three of which (KL25, KL23, and KL122) were present among the 30 most prevalent identified globally as potential vaccine targets in a recent metanalysis of 1930 Kp neonatal blood isolates from 13 countries, with KL25 ranking as the third most prevalent (49). Although our study consisted of only a small sample size, this nevertheless suggests these K loci may be important targets for vaccine development within the community as well as the hospital setting. There is interest in development of Kp vaccines for the prevention of neonatal sepsis through maternal vaccination as well as for other high-risk groups (50). Importantly, however, assuming a duration of effect of a maternal vaccination candidate of three months, only 2 of the 7 infants and children would have been protected by a hypothetical maternal vaccine in this small study of ALRI deaths in the community. We believe our findings call for larger and more systematic studies of Kp causing severe community ALRI in infants and children to inform vaccine target profiles and policy.

Plasmids are major vehicles through which Kp acquires and disseminates ARGs and virulence determinants via horizontal gene transfer. MDR-carrying plasmids can circulate among Kp lineages and even transfer to other species, facilitating ARG dissemination and contributing to AMR spread and difficult-to-control outbreaks (51, 52). In this study, several clinically important plasmid replicons were detected, including FIB(K), IncFII(K), Inc(R), ColRNAI, IncFII(pKP91), IncFIAHI1, IncN, IncFII(pHN7A8), and Col(pHAD28). These replicons are frequently associated with MDR plasmids and high-risk global Kp clones (28, 36, 37). Although detection of the same plasmid replicon across multiple Kp sublineages suggests possible horizontal gene transfer between these lineages, further investigation using long-read sequencing would be needed to confirm this.

Several data points support the validity of the genomic analysis generated through this novel application of RNA-probe hybridization-capture of Kp from lung tissue samples. Kp C_q_ values from the FTD-32 multiplex PCR correlated highly to a singleplex Kp qPCR and to the on-target Kp reads from RNA-probe hybridization-capture sequencing. We observed high concordance in detected SLs, CGs, K-loci, ARGs, and acquired virulence determinants between paired samples obtained from the same deceased individual. Previously identified SL and K-locus associations were detected, including the association between SL280 and KL23, SL607 and KL25, SL17 and KL122, as well as SL37 and KL23 (27, 41, 42, 46). Additionally, SLs from our study that overlapped with strains reported in other recent sub-Saharan African studies in Zambia and Malawi showed highly similar ARG profiles, including acquired aminoglycoside, β-lactamase, fluoroquinolone, sulphonamide, tetracycline, and trimethoprim resistance genes (27, 46). Our results thus demonstrate this method allows detailed analysis of genetic epidemiology including STs (MLST and cgMLST), serotyping, acquired AMR and virulence of Kp lineages from tissue samples. RNA-probe hybridization-capture may be useful for other settings, including post-mortem surveillance studies, where genetic epidemiology investigation is warranted but cultures are not available. The costs, probe-target development needs, and inability to perform phenotypic testing, however, may limit its broader applicability.

This study was not without limitations. First, the small sample size limits the generalizability of the sequencing findings. Larger, representative studies incorporating genetic analyses of Kp causing severe pediatric ALRI in the community are urgently needed. Second, determining the pathogenic cause in ALRI is complex and the sequencing findings in our study do not remove uncertainty from the true cause of death for the infants and children in this study. To incorporate multiple modalities of evidence into this determination, we relied on the collective clinical reasoning of a DeCoDe panel of experts. The biopsies that underwent Kp sequencing in this study had multiple pieces of evidence in support, first, of a respiratory cause of death and, second, of Kp as the causative pathogen. However, some cases contained multiple conflicting pieces of evidence, for example one case (**Table 1**), where the same three virulent pathogens were detected on multiple biopsies, leading the DeCoDe panel to give a designation of moderate certainty for the diagnosis of Kp ALRI.

The RNA-probe hybridization-capture technique also had limitations. High human DNA contamination despite hybridization-capture resulted in low Kp reads in several samples, which limited the resolution of lineage typing and detection of ARGs. While sufficient for analysis, higher coverage would have given confidence to some of the lower coverage hits. In addition, the method may have benefited from a second round of hybridization-capture as with other high sensitivity RNA-probe hybridization-capture approaches. Moreover, while the acquired ARGs and virulence determinants we detected frequently associate with Kp, especially within sub-Saharan Africa, we could not definitively attribute these to Kp over other species potentially present at low abundances within this study. For example, *iuc5* which was detected in case 4803, is also commonly associated with *E. coli*, of which reads were detected in all samples. Although it is possible these *E. coli* reads may have arisen from misclassification during taxonomy profiling, due to shared homology between Kp and *E. coli*, they may also suggest background contamination, possibly due to increased gut barrier permeability post-mortem. Given these uncertainties, future culture-based isolation and whole genome sequencing studies would be required to confirm the association of these genes with the detected Kp lineages.

## Conclusions

This small study characterizing Kp genomes from Zambian children who died of suspected Kp ALRI in the community warrants further study into the prevalence of severe and fatal Kp ALRI and the underlying genomic epidemiology. This study investigated Kp sequenced from preserved lung biopsies in 7 suspected Kp ALRI deaths identified through a post-mortem MITS study of under-5 community respiratory deaths in Lusaka, Zambia. ALRI remains the second most common cause of under-5 mortality globally and growing evidence suggesting Kp as a contributing pathogen raises concerns around current clinical management approaches. Across the samples, we detected several distinct sublineages of global importance, genes encoding acquired resistance to important antimicrobial classes in all cases and acquired virulence factors in two cases. Given the critical implications for prevention and treatment of severe under-5 ALRI in the community, additional, larger scale genomic epidemiology studies should be carried out in similar populations.

## Methods

This study used a target hybridization-capture metagenomic approach to sequence Kp from metagenomic DNA extracted from post-mortem lung biopsies of infants and children who died outside of hospital settings in Lusaka, Zambia. The methods of the original post-mortem study have been previously detailed (15). Briefly, enrollments occurred at the University Teaching Hospital morgue, which serves the city of Lusaka. All next-of-kin of the deceased approached for the study were offered grief counseling, and, if interested in study participation, provided informed consent followed by a verbal autopsy. The body of the deceased was then brought to the autopsy theater, the thorax was cleaned with ethanol and iodine, and six minimally invasive tissue samples (MITS) were obtained from upper, middle, and lower lung zones from the left and right side along the axillary line. Molecular testing and histopathology was performed on specimens from each of the six lung zones sampled. The molecular testing performed was a 32-respiratory pathogen PCR panel (FTD-33 kit, Fast-Track Diagnostics). For histopathology, lung samples were fixed in neutral buffered formalin, followed by paraffin embedding, microtomy, hematoxylin and eosin staining, and finally, reviewed by two experienced pathologists.

A panel adjudication approach modeled after the Child Health and Mortality Prevention Surveillance Determination of Cause of Death (DeCoDe) Process was used to determine the causes of death in this post-mortem study, based on the totality of subjective and objective data. The panel convened for this study included 7 clinicians with infectious diseases and pediatrics experience practicing in Lusaka. The panel met in person and reviewed case dossiers, including representative photographs of the histopathology for each of the deceased, to achieve a consensus determination of whether the death was due to respiratory or non-respiratory causes, the cause of death, and whether the death was preventable or not. For the present sequencing study, we selected 8 cases with the strongest molecular data (at least two biopsies with Kp detected at qPCR cycle thresholds of less than 30) from among the 13 deceased infants and children adjudicated to have died from Kp pneumonia. We attempted to perform target hybridization-capture sequencing on two separate lung biopsy samples from each of these select cases.

### DNA extraction, quality control, and *K. pneumoniae-*specific ZKIR-qPCR analysis

DNA was extracted using the InviSorb® Spin Universal Kit (Invitek Diagnostics, Germany) following the manufacturer’s protocol for tissue biopsy samples in final elution volume of 60 µL DNA concentration. The concentration and quality of the extracted DNA from each of the lung biopsy samples was measured by Quibit 3.0 fluorometer using dsDNA Narrow Range Assay (Thermo Fisher Scientific, Waltham, USA) and Nanodrop 1000 Spectrophotometer (Thermo Fisher Scientific), respectively. All DNA samples which passed quality control assessment underwent analysis using the highly Kp sensitive and specific ZKIR-qPCR assay as previously described (29, 30). Briefly, 2.5 ng of 10 ng/µL DNA (25 ng total) was used as input for each 25 µL qPCR reaction, with reaction mixture and cycling conditions as described (30). All samples were assayed in triplicate using an Applied Biosystems 7500 Real-Time PCR System (Thermo Fisher Scientific), and criteria used for positive Kp detection were as described previously. Kp genome copy number was calculated from the standard curve using 7500 Real-Time PCR Analysis Software v2.3 (Applied Biosystems, Life Technologies, Waltham, USA) as described (30).

Due to the very high detection sensitivity of the RNA-probe target hybridization-capture method employed in this study, a blank negative control was used to account for background/cross-contamination during the hybridization-capture process which consisted of DNA elution buffer only. This blank sample had no detectable DNA by qubit analysis and was negative for Kp by the ZKIR-qPCR.

### Library construction, targeted RNA probe hybridization-capture, and sequencing

Hybridization-capture of Kp from lung biopsy metagenomic samples was performed by SureSelect XT HS2 targeted RNA probe hybridisation-capture (Agilent, Santa Clara, USA). A custom panel of 145,787 120bp RNA probes (total size: 2.635 Mbp) was designed targeting core and acquired genes from the *Klebsiella pneumoniae* pan-genome, including cgMLST, MLST, K-and O- loci, ARGs, virulence loci, and plasmid replicons (**Supp. Table 11**). Probes were designed with at least 90% sequence homology to the reference sequences with 1–2x tiling density, by an in-house developed algorithm by Agilent Technologies. For regions with ambiguous bases, among all possible sequences, representative probes with a 90% base homology threshold were selected. The algorithm reduces the number of relevant baits to ensure cost efficiency. The final bait sets comprised baits (Design ID S3491653 and 61512) of 120bp RNA probes (total size: 2.635 Mbp).

Sequencing library construction and targeted RNA probe hybridization-capture of samples was performed as per the SureSelect XT HS2 DNA Kit Library Preparation (+/- MBC)/Fast-Hyb Target Hybridization-capture/Post-hybridization-capture Pooling Workflow protocol (Agilent; catalog nr G9982A) with modifications based on the Utilization of Agilent SureSelect Target Hybridization-capture for Whole Genome Sequencing of Viruses and Bacteria protocol (Agilent; catalog nr PR7000-2035). Briefly, 150 - 200 ng DNA was used as input for library construction. Samples with <150 ng DNA were bulked with human genomic DNA (Promega, Madison, USA). Following enzymatic fragmentation, end repair and A-tailing, libraries underwent PCR amplification and indexing, followed by purification on AMPure Beads. 1000 ng of DNA library were hybridized with 1:10 diluted probes for 90 mins according to the protocol. After hybridization, a hybridization-capture step using Streptavidin beads, followed by several steps of washing were done according to the protocol. A final post-hybridization-capture PCR was performed with 19 cycles for samples with > 200 Kp genome copies/ng DNA and 21 cycles for samples with < 200 Kp genome copies/ng DNA.

All target enriched samples were sequenced on the Illumina NextSeq 500 platform at 75 bp paired end reads. Processing of FASTQ files and removal of residual human DNA was performed as previously described (30).

### Analysis of Target-enriched samples

#### Taxonomy profiling

Taxonomic profiling, including total bacterial read count as well as Kp read count and relative abundance estimates were performed using Kraken2 v2.1.2 and Bracken v2.6.1 with MiniKrakenDB_8GB v202003 (53, 54). Quantification of unique Kp and *E. coli* reads in samples was performed using Centrifuge v1.0.4 using the standard p_compressed+h+v database (55).

#### cgMLST and MLST analysis

Kp lineage analysis was performed using both the previously described 7-gene multi-locus sequence typing (MLST) scheme and the core genome multi-locus sequence typing (cgMLST) scheme consisting of 629 core genome genes (32, 33). Both MLST and cgMLST alleles were identified from sequenced reads using the aligner KMA v1.4.18 (56) against either the MLST or cgMLST allelic nucleotide sequence databases downloaded from BIGsdb (https://bigsdb.pasteur.fr/klebsiella/). Detected MLST/cgMLST alleles were filtered to include only those with ≥80% gene coverage and ≥90% sequence identity. The allele corresponding to each MLST/cgMLST gene detected above this threshold that had the highest ConClave score (accumulated alignment score) was then used to construct the final Kp MLST/cgMLST profile for each sample. The resulting detected MLST/cgMLST profiles were compared to the Kp MLST/cgMLST allelic profile database available at BIGSdb to identify the closest matching ST/cgST lineage.

#### K-locus prediction

Within our custom RNA-probe panel were probes targeting the K-locus genes *wzi*, *wzx*, and *wzy*. To predict the Kp K-locus from our samples we used KMA v1.4.18 (56) to align sequence reads to the K-locus gene nucleotide sequence database downloaded from Kaptive v3.0.0 (34) using the command ‘kaptive extract kpsc_k --ffn’. Detected alleles were filtered to include only those with ≥80% gene coverage and ≥90% sequence identity. The allele for each K-locus gene with the highest ConClave score was then retained. K-locus was called if at least two out of the three detected K-locus genes (*wzi*, *wzx*, and *wzy*) corresponded to the same K-locus.

#### Detection of acquired ARGs, virulence loci, and plasmid replicons

Detection of ARGs, virulence loci, and plasmid replicons was performed using the read alignment tool KMA v1.4.18 (56). For ARGs, reads were aligned to the CARD v3.0.8 database. For virulence loci, allelic sequences were downloaded from BIGSdb (https://bigsdb.pasteur.fr/klebsiella/) and ST profiles for yersinabactin (ybt), colibactin (clb), aerobactin (iuc), salmochelin (iro) and rmpADC were analyzed using schemes described previously (57–59). For detection of plasmid replicon sequences, the PlasmidFinder database was used. Presence of a gene was called if it was detected with ≥ 90% sequence identity, ≥ 80% gene coverage, and ≥ 50x coverage depth. Detection of an acquired virulence loci was called if ≥ 50% of the expected alleles for that loci were detected above the minimum thresholds. Lineage was reported if ≥ 50% of the detected alleles matched with an allelic profile when queried against the BIGSdb database.

#### Contig-based analysis

Paired-end reads were assembled into contigs using MegaHit v1.2.9 and --presets meta-sensitive parameter. Assembled contigs were analyzed for ST, acquired virulence loci and ARGs via Kleborate v3.1.2 (19) and capsule and O-antigen loci via Kaptive v3.0.0 (34) using default parameters.

#### Statistical analysis and data visualization

Statistical analysis and data visualization was performed in R v4.5.0 using RStudio 2025.05.01+513.

#### Computational resources

Bioinformatic analysis of target enriched metagenomic samples was performed on the Norwegian academic high-performance computing and storage services maintained by Sigma2 Norwegian Research Infrastructure (NRIS) (http://www.sigma2.no/) (Sigma2 project number: nn9794k).

## Data availability

Raw Illumina sequence reads from target enriched samples (human DNA removed) are available on ENA, Bioproject PRJEB105599.

## Ethics statement

Ethical oversight for the ZPRIME and LCIS studies were provided by institutional review boards at Boston University and the University of Zambia. Written informed consent was obtained from the deceased’s family members or representatives.

## Acknowledgements

The authors thank Sylvain Brisse (Institut Pasteur, France), Kelly Wyres (Monash University, Australia), Erik Hjerde (UiT The Arctic University of Norway), and Ryan Wick (The University of Melbourne, Australia) for assistance in RNA probe target selection and design. We thank Even Østli and his team Agilent for their assistance in RNA-probe production and technical support with the hybridization-capture. We thank Ruth Paulsson and Hagar Taman (the Genome Sequencing Support Centre, Norway) for their assistance with library construction and sequencing. We also thank Tom Stanton (Monash University, Australia) for his advice when analysing Kp K-loci.

## Author contributions

The study was conceptualized by CG, AS, KH, AW, KL, and MS. Funding acquisition was performed by CG, AS, and KL. Participant recruitment, grief counseling, and sample acquisition were performed by BN, CC, SC, LP, DN, and JL. Adjudications were performed by NN, EJM, MK, CM, BM, LN, VM, SWS, and coordinated by MC. Lung biopsy DNA sample preparation and shipping was organized by AW and MS. RNA-probe synthesis was coordinated by KL and AW. RNA-probe hybridization capture of samples was performed by WK. DNA sequencing and library preparation was arranged by KL and WK. Data analysis was performed by KL with clinical summaries compiled by AW. KH and AS provided expertise on *Klebsiella pneumoniae* genomics and clinical aspects. AW and KL prepared the first manuscript draft. All authorships contributed with reviewing and editing the manuscript and approved the final version.

## Abbreviations

ALRI: acute lower respiratory infection
AMR: antimicrobial resistance
ARG: antimicrobial resistance gene
CHAMPS: Child Health and Mortality Prevention Surveillance
CG: clonal group
cgMLST: core genome multilocus sequence typing
cgST: core genome sequence type
ESBL: Extended spectrum beta-lactamase
Kp: *Klebsiella pneumoniae*
LMIC: low- to- middle-income countries
MDR: multidrug resistant
MITS: minimally invasive tissue sampling
MLST: multilocus sequence typing
SL: sublineage
ST: sequence type

