## Supplementary Figures for "Genomic analysis of *Klebsiella pneumoniae* causing community-acquired respiratory deaths among Zambian infants and children using targeted RNA-probe hybridization-capture metagenomics"

**(short title) *Klebsiella pneumoniae* fatal community-acquired pneumonia in Zambian children**

**Supplementary Figures**

Kenneth Lindstedt^1,2,3,^*, Alyse Wheelock^4,^*, Mulemba Samutela^5^, Wasifa Kabir^6^, Mwelwa Chasaya^7^, Natasha Namuziya^8^, Emilia Jumbe Marsden^9^, Monica Kapasa^8^, Chibamba Mumba^10^, Bwalya Mulenga^10^, Lisa Nkole^8^, Rachel Pieciak^11^, Victor Mudenda^10^, Chilufya Chikoti^7^, Benard Ngoma^7^, Charles Chimoga^7^, Sarah Chirwa^7^, Lilian Pemba^7^, Diana Nzara^7^, James Lungu^7^, Leah Forman^12^, Edgar Simulundu^13, 14^, William MacLeod^11^, Crispin Moyo^7^, Somwe Wa Somwe^5,8^, Kathryn E. Holt^15,16^, Arnfinn Sundsfjord^1,2,3^, Christopher J. Gill^11^

1 Norwegian Centre for Detection of Antimicrobial Resistance, Department of Microbiology and Infection Control, University Hospital of North Norway, Norway

2 Department of Medical Biology, Faculty of Health Sciences, UiT the Arctic University of Norway, Norway

3 Centre for New Antibacterial Strategies, UiT the Arctic University of Norway, Norway

4 Section of Preventive Medicine and Epidemiology, Chobanian & Avedisian School of Medicine, Boston University, Boston, MA, USA

5 School of Health Sciences, University of Zambia, Lusaka, Zambia

6 Aquaculture and Environment Group, UiT the Arctic University of Norway, Norway

7 Avencion Limited, Lusaka, Zambia

8 University Teaching Hospital – Children’s Hospital, Lusaka, Zambia

9 Pendleton Children’s Clinic, Lusaka, Zambia

10 Department of Pathology, University Teaching Hospital, Lusaka, Zambia

11 Department of Global Health, Boston University School of Public Health, Boston, MA, USA

12 Center for Health Data Science (CHDS), Boston University School of Public Health, Boston, MA, USA

13 Macha Research Trust, Choma, Zambia

14 Zambia National Public Health Institute, Lusaka, Zambia

15 Department of Infection Biology, London School of Hygiene and Tropical Medicine, London, UK

16 Department of Infectious Diseases, School of Translational Medicine, Monash University, Melbourne, Victoria 3004, Australia

*Co-first authors, contributed equally

Corresponding author (technical questions): Kenneth Lindstedt, MBBS, PhD, Norwegian Centre for Detection of Antimicrobial Resistance, Department of Microbiology and Infection Control, University Hospital of North Norway, Norway.

Corresponding author (study information): Alyse Wheelock, MD, Section of Preventive Medicine and Epidemiology, Chobanian & Avedisian School of Medicine, Boston University, Boston, MA, USA.

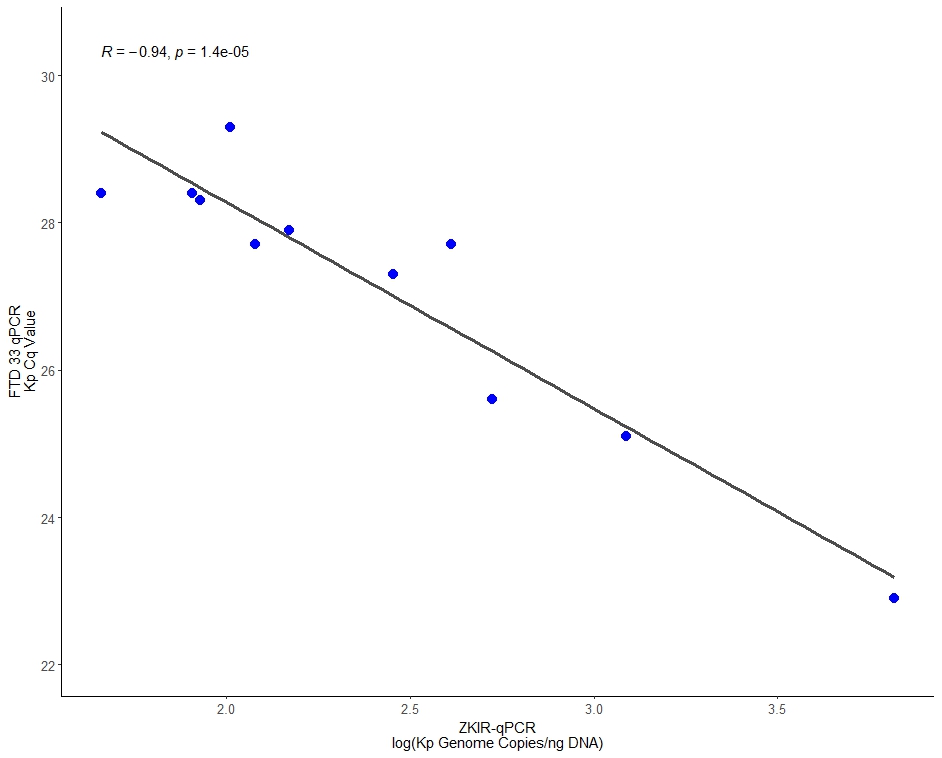

**Supplementary Figure 1.** Comparison of Kp detection from the 11 lung tissue DNA samples from 7 deceased infants and children by the FTD-33 multiplex qPCR and the ZKIR-qPCR. *R* = Pearson’s correlation coefficient.

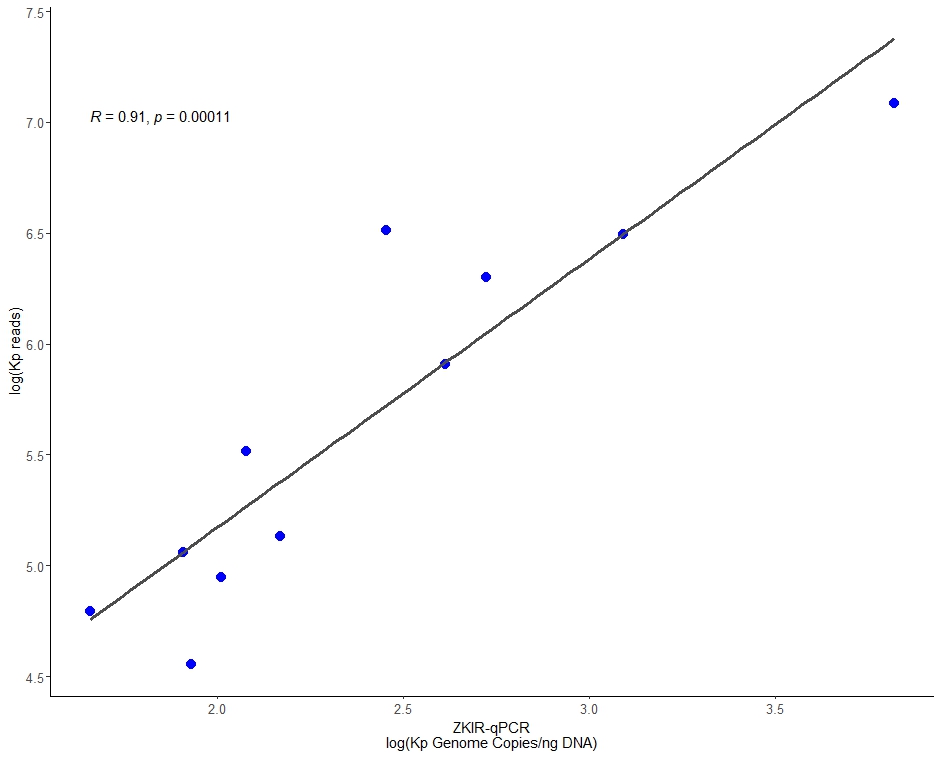

**Supplementary Figure 2**. Comparison of reads assigned to Kp in samples post-capture (determined by the taxonomic profiler Kraken2 + Bracken) vs. abundance of Kp in pre-capture samples (measured by the ZKIR-qPCR). *R* = Pearson’s correlation coefficient.

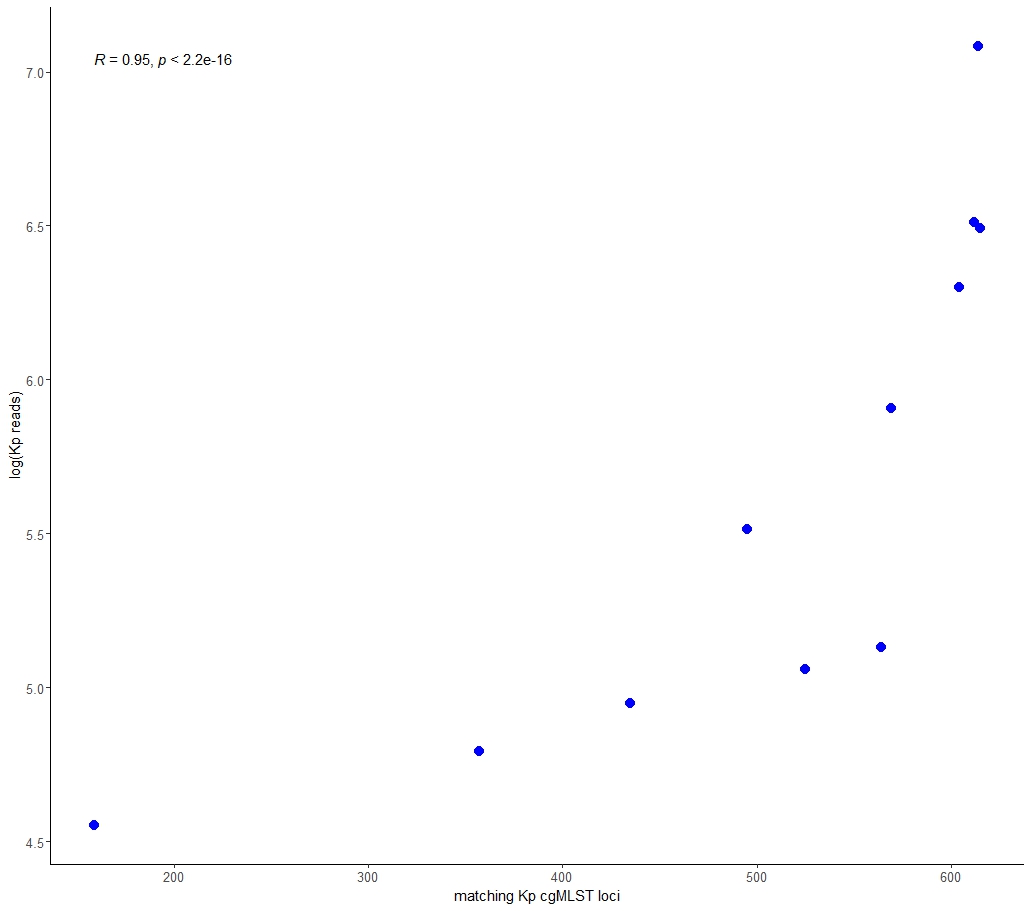

**Supplementary Figure 3**. Comparison of detected Kp cgMLST loci that matched to alleles of a cgMLST profile in the BIGSdb Kp cgMLST database vs. total reads assigned to Kp post-enrichment. *R* = Spearman’s rho.

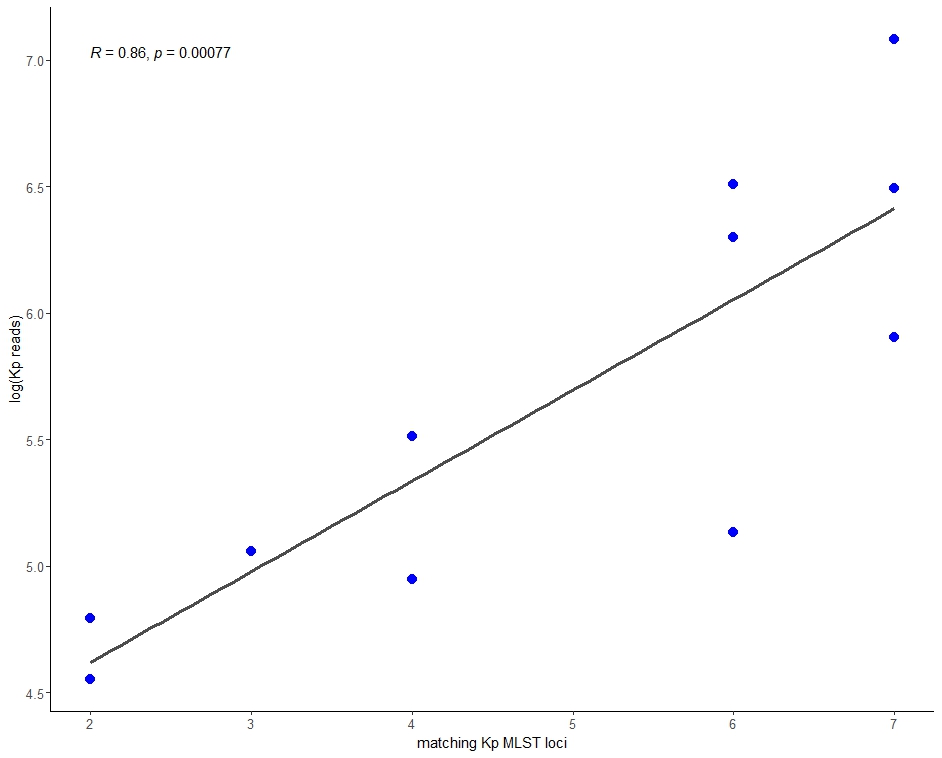

**Supplementary Figure 4.** Comparison of detected MLST loci (out of 7) that matched to the expected MLST profile based on our cgMLST analysis vs. total reads assigned to Kp. *R* = Spearman’s rho.
